# Macrophages in the uterus are functionally specialised and continually replenished from the circulation

**DOI:** 10.1101/2021.09.02.458731

**Authors:** Nicholas A. Scott, Lamiya Mohiyiddeen, Livia Lacerda Mariano, Peter T. Ruane, John D. Aplin, Olivia C. Moran, Daniel R. Brison, Molly A. Ingersoll, Elizabeth R. Mann

**Author notes:** Equal contribution (Nicholas A. Scott, Lamiya Mohiyiddeen, Livia Lacerda Mariano).

## Abstract

Macrophages are innate immune cells that fight infection but also regulate tissue regeneration and remodelling. In the uterus, although tissue remodelling is essential for establishment and maintenance of pregnancy, the specialisation of macrophages is not well characterised compared to other mucosal tissues. Here we show that uterine macrophages are functionally specialised, expressing multiple markers of alternative activation associated with tissue remodelling and repair, and responding more highly to the type 2 cytokine IL-4 than other mucosal tissue macrophages. Uterine macrophages were continuously replenished from circulating bone marrow-derived CCR2^+^ monocytes that fluctuated dramatically in number throughout the reproductive cycle, and had properties distinct from the macrophages that they became, including differential responses to microbial stimulation. Importantly, many of these properties of uterine monocytes and macrophages were conserved between mice and humans. These findings further our understanding of immune regulation of uterine tissue integrity and have important implications for differences in immune responses to infections at different phases of the reproductive cycle.

**SUMMARY:** Uterine macrophages are specialised, alternatively activated cells that are replenished from circulating bone marrow-derived monocytes. Monocyte and macrophage properties fluctuate markedly throughout the reproductive cycle, with many features conserved between mice and humans, and exhibiting differential responses to microbial stimulation.

## INTRODUCTION

The female reproductive tract faces unique challenges, particularly in the uterus lining (endometrium) which undergoes repeated shedding, regeneration and remodelling during the menstrual cycle^1,2^. This allows embryo implantation to occur, while disturbances can manifest as miscarriage and poor pregnancy outcomes. Macrophages are innate immune cells that are not only the first line of defence against invading pathogens, but also play key roles in tissue homeostasis, including remodelling, regeneration and repair^3^. These processes are essential for endometrial function^4^ and tight regulation is critical, as inappropriate or prolonged macrophage activation can lead to chronic inflammation, structural defects and scarring (fibrosis)^3^. Thus, macrophages are proposed to contribute to regulating tissue integrity in the uterus and localise to areas of tissue repair and remodelling in a mouse model of menstruation^5^.

Classical activation of macrophages is usually associated with pro-inflammatory immune responses and production of additional cytokines such as TNFα, leading to effector cell responses and killing of intra-cellular micro-organisms or tumour cells. By contrast, alternatively activated macrophages are typically polarized by exposure to type 2 stimuli, including cytokines IL-4 and IL-13, and promote tissue remodelling, repair and angiogenesis. Alternatively activated macrophages are characterised by expression of scavenger receptors including CD163 and the mannose receptor CD206, the antigen presentation molecule MHC class II and resistin-like molecule α (Relmα)^6–8^. *In vivo*, tissue macrophages are a highly heterogenous group of cells that exist in a range of different states^9^, which incorporate varying degrees of classical and alternative activation features. Nonetheless, these polarization phenotypes can be found in environments with inflammatory pathologies (classically activated macrophages) or wound healing and tissue repair environments (alternatively activated macrophages)^10,11^.

Despite a wealth of information on macrophages from many different organs, macrophages in the uterus have not been characterised in anywhere near the depth as macrophages from other mucosal sites such as the intestine, lungs and bladder, where they are tailored for their tissue-specific functions^12–14^. This limited level of insight into immune mechanisms in the female reproductive tract means there is little information regarding the ontogeny, properties and functions of macrophages in the uterus, and thus a deficit in understanding of infertility and miscarriage pathology.

Here, we show that macrophages have distinct characteristics of alternative activation in the uterus and are distinct from macrophages in other mucosal tissues due to their potent responsiveness to type 2 cytokines, such as IL-4. Macrophages in the uterus are predominantly replenished from circulating monocytes with functional differences between recruited versus tissue-resident macrophages related to alternative activation. Importantly, many properties of monocytes and macrophages in the uterus are conserved between mice and humans, in particular, the marked increase in numbers of monocytes following ovulation and fluctuations in macrophage proliferation throughout the reproductive cycle. These alterations in monocyte and macrophage dynamics in the uterus have implications not just for changes in tissue remodelling capacity, but also for immune responses to infection within the reproductive tract.

## RESULTS AND DISCUSSION

### Murine uterine macrophages are highly responsive to the type 2 cytokine IL-4

In some tissues, macrophages are differentiated from monocytes migrating from the bone marrow and peripheral blood during homeostasis and inflammation^15,16^. It is important to distinguish monocytes from macrophages in tissues given the striking differences in their function. For example, recruited monocytes in the intestine are potently responsive to stimulation with the bacterial cell wall component lipopolysaccharide (LPS) whilst a hallmark feature of gut macrophages differentiated from recruited monocytes is their hyporesponsiveness to bacterial stimulation^17^. Here, we identified monocytes and macrophages in the murine uterus from a heterogenous pool of CD11b^+^CD64^+^ cells selectively expressing the monocyte marker Ly6C and macrophage-associated protein F4/80 (Fig. 1A). F4/80^+^ macrophages were distributed throughout the mouse endometrium, forming a dense network around the regularly regenerated luminal epithelium (Fig. 1B).

**Figure 1:**
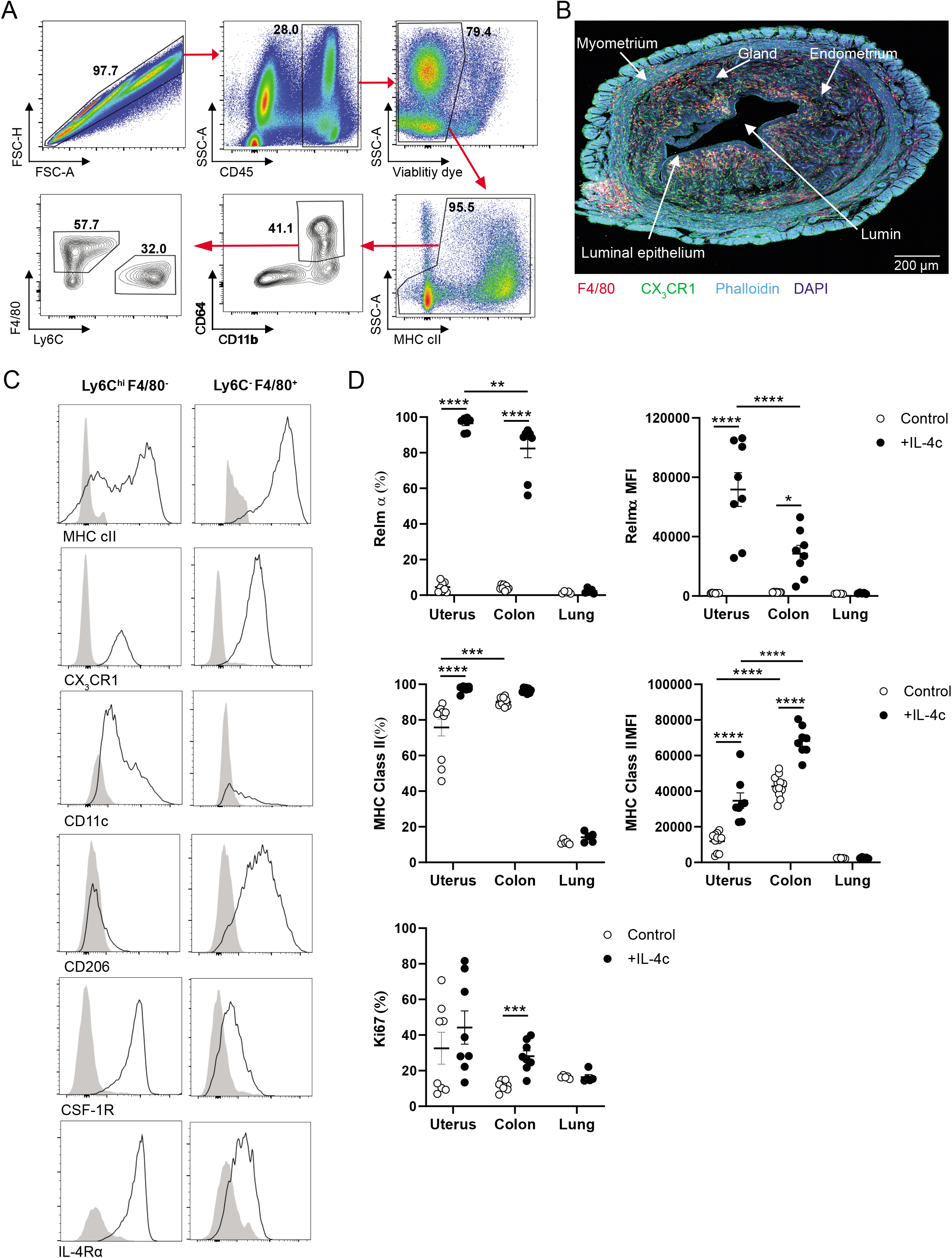
Characterisation of uterine macrophages in mice. **(A)** Representative flow cytometry plots depicting gating strategy for identification of monocytes and macrophages in the uterus in 8-10 week old C57BL/6 mice. **(B)** Representative confocal image of a uterus from a CX3CR1^GFP^ C57BL/6 mouse at 20x, merged image. **(C)** Histograms show the relative expression of MHC class II, CX3CR1, CD11c, CD206, CSF-1R and IL-4Rα on uterine monocyte and macrophage subsets. Negative control staining shown as filled grey histograms on each plot (fluorescence minus one; FMO). Histograms representative of >5 independent experiments. **(D)** C57BL/6 mice were injected with PBS or IL-4c i.p. on day 0 and day 2, and uterine, colonic and lung tissue collected on day 4. Experiments were repeated for alveolar macrophages using i.n. administration of IL-4c on day 0 and day 2, and lung tissue collected on day 4. Expression of Relmα, MHC class II and Ki67 was assessed by flow cytometry from single cell suspensions following tissue digest in all cases. Summary data shown are representative of 2 independent experiments (MHC class II: n=12 for uterus and colon, n=5 for lung (i.n.); Relmα and Ki67: n=8 for uterus and colon, n=5 for lung (i.n.). Data shown as mean ± SEM and statistical comparisons were performed using a 2-way ANOVA with Tukey post-hoc corrections. For all experiments, **p<0.01; ***p<0.001; ****p<0.0001.

All F4/80^+^ macrophages expressed MHC class II, whereas Ly6C^hi^ monocytes existed as two populations of MHC class II^+^ and MHC class II^-^ cells (Fig. 1C). These data suggest that monocytes and macrophages in the uterus may be capable of local antigen presentation to T cells. Uterine monocytes and macrophages uniformly expressed the fractalkine receptor CX3CR1 (Fig. 1B and 1C), a key regulator of macrophage function at sites of inflammation^18,19^, with CX3CR1 knockout mice displaying impaired cutaneous wound healing^18^. Taken with evidence that uterine-derived fractalkine can induce properties of alternative activation in macrophages^20^, the presence of CX3CR1 on uterine monocytes and macrophages supports a role for these cells in healing and repair processes. Accordingly, uterine macrophages expressed low levels of CD11c which is associated with pro-inflammatory, classically activated macrophages^21^. Nearly all macrophages expressed the macrophage mannose receptor CD206, one of the hallmarks of alternatively activation^6^, although this protein was largely absent from monocytes. Monocytes and macrophages in the uterus expressed differential levels of CSF-1R, which is required for monocyte differentiation into macrophages and, in particular, the development of macrophages that regulate tissue structure^22^. Both populations also expressed IL-4Rα, to different extents, suggesting that they are responsive to IL-4-mediated alternative activation of macrophages^23^ (Fig. 1C).

We investigated whether uterine macrophages were functionally similar to macrophages in the mucosal tissues of the gut and the lung following systemic administration of recombinant IL-4 complexed with mAb to IL-4 (IL-4c), which extends the bioactive half-life of the cytokine^24^. Uterine macrophages demonstrated a marked response to IL-4, with potent induction of Relmα, a functional signature of alternatively activated macrophages^7^. IL-4 induced a higher proportion of Relmα^+^ macrophages in the uterus alongside higher levels of expression of Relmα in comparison with macrophages from the intestine or alveolar macrophages in the lung (Fig. 1D). IL-4 also increased levels of MHC class II on uterine macrophages. IL-4 induced proliferation of macrophages in the colon as suggested by elevation of the cell-cycle marker Ki67. In uterine macrophages, Ki67 expression was extremely variable, ranging from approximately 10-80% of macrophages (Fig. 1D).

These data indicate that monocytes and macrophages are abundant throughout the endometrium, expressing features indicative of an involvement in tissue repair and remodelling. The potent response of uterine macrophages to the type 2 cytokine IL-4 indicates that these cells are readily polarised towards alternative activation, further supporting a role for uterine macrophages in maintenance of tissue integrity.

### Dynamic changes in monocytes and macrophages in the uterus throughout the reproductive cycle in mice

We hypothesized that the extensive variability of macrophage expression of Ki67 in the uterus may be due to the cyclical changes in local hormones. Indeed, changes in macrophage numbers are reported in the uterus of mice and rats throughout their reproductive (estrous) cycle^25–27^. However, the phases of the cycle where macrophages numbers are highest vary among studies with results potentially confounded by differences among species and strains, specific localization, and the use of molecules to identify macrophages such as OX-41/CD172a^27^, which are expressed by other myeloid cells.

We, therefore, staged mice at the time of tissue collection as being in one of the four estrous cycle stages of metestrus, diestrus, proestrus or estrus^28^. We first assessed the total proportions and numbers of Ly6C^hi^ monocytes and F4/80^+^ macrophages from whole tissue digests and characterisation by flow cytometry as above (Fig. 1). Macrophages ranged from 10-30% of total uterine CD45^+^ leucocytes but no significant differences were observed in proportions or numbers of macrophages in the uterus among the different phases of the estrous cycle. By contrast, a marked increase in proportions and numbers of monocytes in the uterus were observed during estrus, directly following ovulation when estrogen levels decrease. In some cases, monocytes comprised over 10% of all CD45^+^ leucocytes during estrus (Fig. 2A). These data indicate the importance of assessing monocytes and macrophages separately in the uterus, despite shared expression of several cell surface molecules used to identify them.

**Figure 2:**
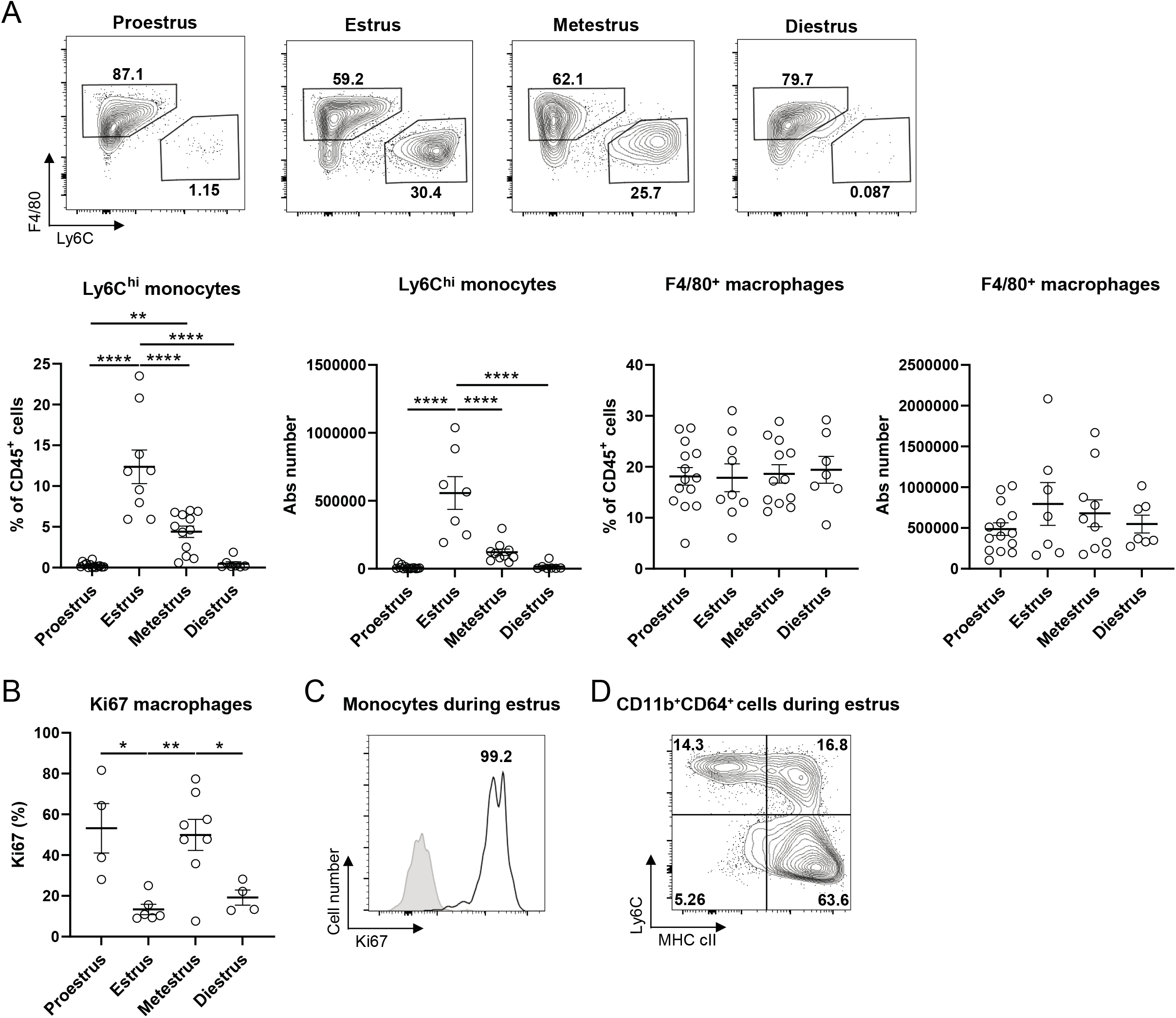
Dynamics of uterine monocytes and macrophages during the murine estrous cycle. **(A)** Representative flow cytometry plots demonstrating uterine monocytes and macrophages from 8-10 week old C57BL/6 mice during the four stages of the estrous cycle; proestrus, estrus, metestrus, diestrus. Summary data showing of proportions of total CD45^+^ cells and absolute numbers are representative of >5 independent experiments (monocyte and macrophage proportions of CD45^+^ cells: proestrus n=14, estrus n=9, metestrus n=12, diestrus n=7; monocyte and macrophage absolute numbers: proestrus n=14, estrus n=7, metestrus n=10, diestrus n=7). Data shown as mean ± SEM and statistical comparisons were performed using 1-way ANOVA with Tukey post hoc corrections. **(B)** Summary data showing proportions of uterine macrophages expressing Ki67 (proestrus n=4; estrus n=6; metestrus n=8; diestrus n=4). Data are representative of 3 independent experiments, shown as mean ± SEM and statistical comparisons were performed using 1-way ANOVA with Tukey post hoc corrections. **(C)** Representative histogram demonstrating monocyte expression of Ki67 during estrus; representative of 3 independent experiments. Negative control staining shown as filled grey histogram (FMO). **(D)** Representative flow cytometry plot demonstrating expression of MHC class II and Ly6C on heterogenous pool of monocytes and macrophages (CD11b^+^CD64^+^ cells) during estrus. Representative of 4 independent experiments. For all experiments: *p<0.05; **p<0.01; ***p<0.001; ****p<0.0001.

To assess whether the variability observed in Ki67 expression by uterine macrophages (Fig. 1D) may be caused by differential expression of Ki67 throughout the estrous cycle, we next assessed Ki67 expression in macrophages at different stages of the cycle. Indeed, we revealed striking differences in macrophage Ki67 expression among cycle stages, which fluctuated between high, variable levels of Ki67 during proestrus and metestrus but consistently low levels during estrus and diestrus (Fig. 2B). It was not possible to assess Ki67 expression in monocytes at different stages of the estrous cycle due to the limited numbers or absence of monocytes during proestrus and diestrus (Fig. 2A). Thus, Ki67 expression was assessed in monocytes during estrus when numbers were high. We observed that all monocytes expressed Ki67, suggestive of monocyte proliferation (Fig. 2C). Although these data could indicate that the striking increase in monocyte numbers during estrus may be at least in part due to local proliferation as shown in other tissues^29^, assessment of the heterogenous pool of CD11b^+^CD64^+^ monocytes and macrophages during estrus revealed the presence of a population of MHC class II^+^ Ly6C^+^ monocytes (Fig. 2D). In other mucosal tissues, this population represents an intermediate population of monocytes in a differentiation continuum from incoming Ly6C^+^ monocytes to Ly6C^-^ MHC class II^+^ macrophages^17^. Thus, it is plausible that some monocytes, whether recruited or expanded by local proliferation, may differentiate to maintain the uterine macrophage pool.

### Uterine macrophages are predominantly replenished by circulating monocytes

Macrophages in the uterus expressed varying levels of cell-cycle marker Ki67 throughout the estrous cycle (Fig. 2B), suggesting that some macrophages may be in cell cycle and therefore proliferating to varying degrees through the cycle. Differences in proliferation between macrophages subsets have been previously demonstrated in other tissues e.g. higher levels of proliferation from recently recruited macrophages compared to long term resident macrophages in the peritoneal cavity^30^. Macrophage origins can have important implications for their function depending on the context; for example, recently recruited macrophages can be critical for regeneration in the liver^31^ and the lung^32^, whereas tissue resident macrophages, derived from embryonic precursors, promote tissue repair in the heart following myocardial infarction^33^. Recruited versus resident macrophage subsets also have differential responses to bacterial infections in the bladder^14^ and recruited macrophages can exhibit pro-inflammatory properties to play a key role in disorders involving dysregulation of tissue structure, such as fibrosis^3,34^, or immunosuppressive properties during helminth infection^35^.

To determine whether macrophages in the uterus are maintained by homeostatic replenishment from circulating bone marrow-derived cells, we used shielded irradiated chimeric mice, in which adult animals are irradiated with a lead cover over the uterus. This procedure protects the uterus from radiation-induced immune cell death and the generation of an empty immune cell niche for immune cell infiltration under non-homeostatic conditions. Irradiated CD45.1 animals were transplanted with congenic CD45.2 bone marrow to differentiate immune cell populations resident in the uterus from infiltrating bone marrow-derived cells over time. In these experiments, we compared the level of chimerism in uterine macrophages to that of colonic macrophages from the same animal (with the lead cover also shielding the intestines and protecting from radiation-induced immune cell death), as colonic macrophages are predominantly replenished from the circulation^16,36^.

By 8 weeks, approximately 40% of macrophages in the uterus were replenished by circulating cells whilst colonic macrophages were already predominantly comprised of cells from the circulation. By 14 weeks post-irradiation, the turnover of macrophages in the uterus was comparable to that of macrophages in the colon, which was also evidenced at 20 weeks (Fig. 3A). These results reveal that macrophages in the uterus of adult female mice are predominantly replenished by circulating bone marrow-derived cells. This occurs at a slower rate than colonic macrophages but by 14 weeks the uterine macrophage population is comprised of recruited monocyte-derived macrophages with one of highest replenishment rates of all tissues. Given the high uniform levels of Ki67 in uterine monocytes (Fig. 2C), the varying levels of Ki67 expression by uterine macrophages (Ly6C^-^ F4/80^+^) may be representative of recently recruited cells retaining Ki67 that have rapidly lost Ly6C as seen in other tissues^37,38^. The continual replenishment of macrophages by circulating cells in the intestine is proposed to be in part due to the ongoing exposure to bacterial stimulation from the trillions of commensal bacteria in the gut^36^. However, in the uterus, given a relatively limited exposure to bacterial species, it is possible that the cells are continually recruited and differentiate into macrophages due to the dynamic structural environment and differential requirements for tissue remodelling^4^.

**Figure 3:**
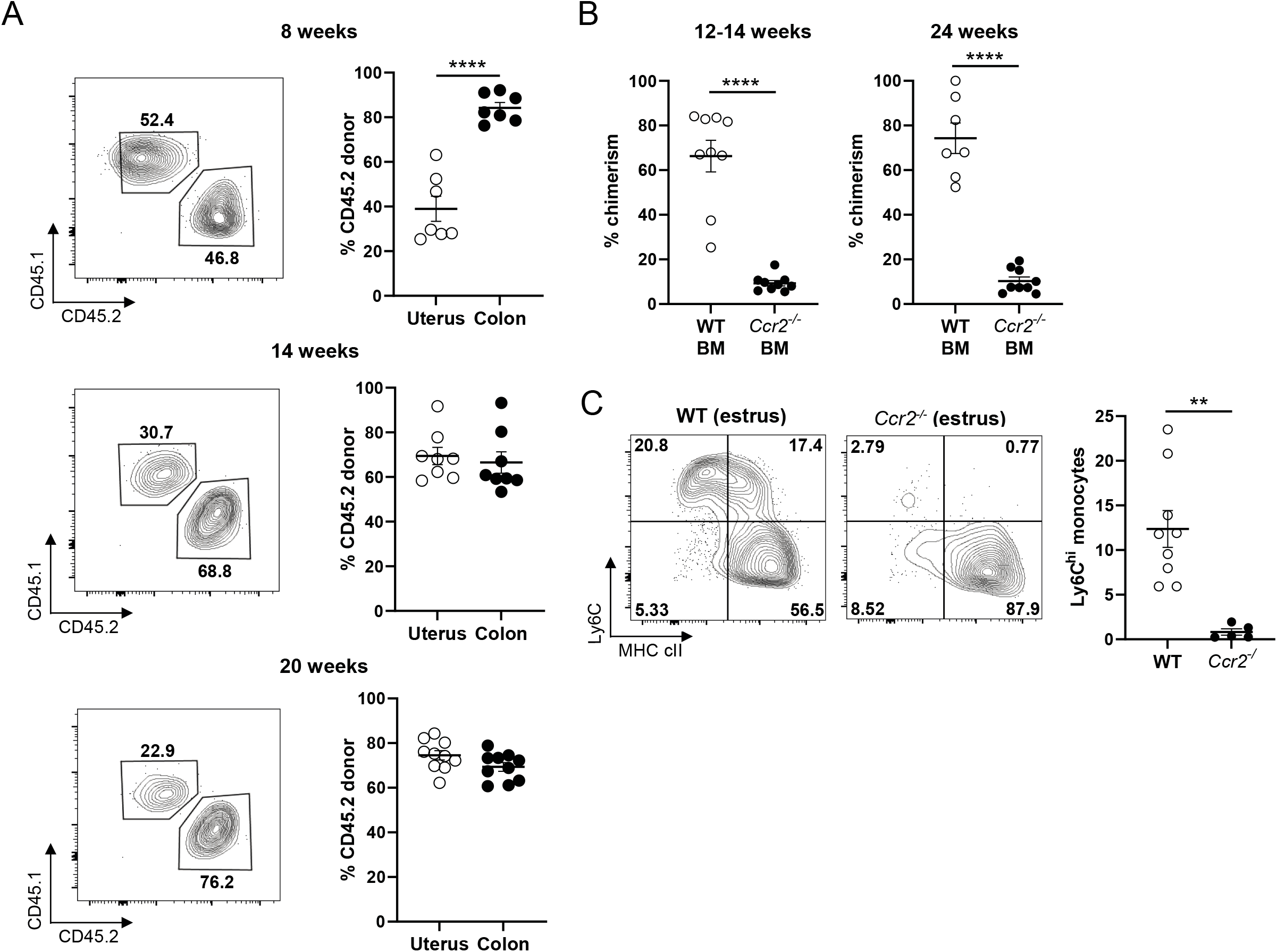
Uterine macrophages in mice are replenished from circulating CCR2^+^ bone-marrow derived cells. **(A)** Shield-irradiated CD45.1 C57BL/6 mice reconstituted with CD45.2 bone marrow and representative flow cytometry plots showing proportions of CD45.1 host and CD45.2 donor cells within the CD11b^+^CD64^+^F4/80^+^Ly6C^-^ macrophage pool in the uterus. Summary data show percentages of CD45.2 donor cells of the total CD11b^+^CD64^+^F4/80^+^Ly6C^-^ macrophage pool in the uterus and colons of the same animals at 8, 14 and 20 weeks. Chimerism of tissue macrophages was normalised to that of circulating monocytes. Data are representative of 4 independent experiments (8 weeks: n=7; 14 weeks: n=8; 20 weeks: n=10), shown as mean ± SEM, and statistical comparisons were performed using paired *t*-tests. **(B)** Shield-irradiated CD45.2 C57BL/6 mice were reconstituted with *Ccr2*^+/+^ CD45.1 bone marrow (labelled as WT BM; n=9 at 12-14 weeks; n=7 at 24 weeks) and shield-irradiated CD45.1 C57BL/6 mice were reconstituted with CD45.2 *Ccr2^-/-^* BM (labelled as *Ccr2^-/-^* BM; n=9 at 12-14 weeks; n=9 at 24 weeks). Chimerism of uterine macrophages was normalised to that of circulating eosinophils, since this was the only circulating immune cell population that demonstrated comparable levels of chimerism in mice reconstituted with WT or *Ccr2^-/-^* BM (S.Fig.3). Data are representative of 2 independent experiments, shown as mean ± SEM and statistical comparisons were performed using unpaired *t*-tests. **(C)** Non-chimeric WT and *Ccr2^-/-^* mice were assessed for monocytes in the uterus during estrus. Representative flow cytometry plots show expression of Ly6C and MHC class II from CD11b^+^CD64^+^ cells, comprising monocytes and macrophages. Summary graph is representative of 2 independent experiments (control: n=9; *Ccr2^-/-^*: n=5). Data are shown as mean ± SEM and statistical comparisons were performed using unpaired *t*-tests. For all experiments, **p<0.01; ****p<0.0001.

To determine whether recruited macrophages were derived from circulating CCR2^+^ monocytes as in other tissues^16^ we also carried out experiments in which shielded irradiated mice were transplanted with *Ccr2*^-/-^ or wild-type (WT) bone marrow (BM). Chimerism in uterine macrophages was markedly reduced in *Ccr2*^-/-^ BM recipients with virtually no replenishment of macrophages from circulating cells, indicating that CCR2^+^ monocytes replace uterine macrophages (Fig. 3B). In further support of a requirement of CCR2 for monocyte entry into the uterus, we assessed monocyte levels in the uterus of *Ccr2*^-/-^ mice during the estrus phase when WT mice have a striking increase in monocyte numbers (Fig. 2B). Ly6C^+^ monocytes in the uterus were nearly completely absent in non-chimeric untreated *Ccr2*^-/-^ mice during estrus, indicating monocyte infiltration to the uterus is dependent on CCR2 (Fig. 3C). Given the large fluctuations in monocyte numbers throughout the estrous cycle but relative stability in macrophage numbers (Fig. 2B), it is likely that not all monocytes differentiate into macrophages. Indeed, the estrous cycle lasts 4 days whereas it takes up to 14 weeks for uterine macrophages to become replenished from circulating monocytes (Fig. 3A). It is possible that some monocytes may differentiate into other cell types, such as dendritic cells, or have a relatively short life span with high levels of apoptosis. Accordingly, there are high levels of co-localisation of cleaved caspase 3 (indicating apoptosis) within CSF-1R-expressing cells in the uterus in a mouse model of menstruation, consistent with programmed cell death of monocytes following completion of endometrial repair^5^.

### Circulating monocyte-derived macrophages in the uterus exhibit enhanced properties of alternative activation compared to their tissue resident counterparts

We demonstrated that uterine macrophages exhibit various properties associated with alternative activation (Fig. 1). Alternatively activated macrophages can be derived from either proliferation of tissue-resident macrophages or those differentiated from recruited monocytes, but these populations are functionally distinct^39^. To determine whether there are differences between recruited versus tissue resident macrophage populations, we compared these populations using our congenic shield-irradiated chimeric mice. Recruited macrophages arising from donor circulating monocytes expressed higher levels of the mannose receptor CD206 than their tissue resident (host derived) counterparts (Fig. 4A), with CD206 being one of the hallmark signatures of alternative activation. However, a greater proportion of tissue resident macrophages expressed Ki67, in keeping with maintenance of this population by local self-renewal (Fig. 4B).

**Figure 4:**
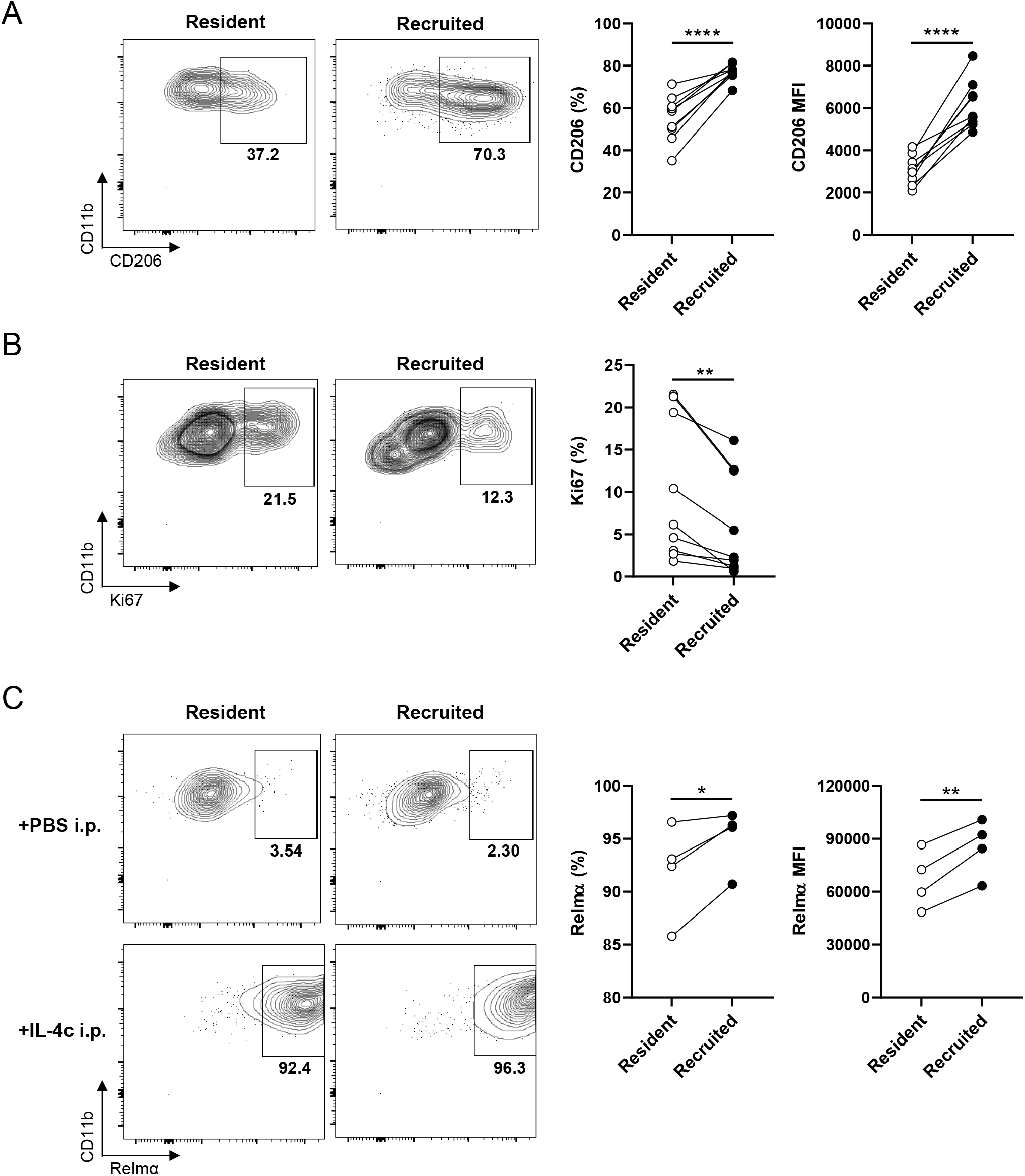
Differential properties of recruited versus resident macrophages in the uterus. **(A)** Shield-irradiated CD45.1 C57BL/6 mice reconstituted with CD45.2 bone marrow and representative flow cytometry plots showing proportions of CD45.1 host (resident) and CD45.2 donor (recruited) macrophages (gated as CD11b^+^CD64^+^F4/80^+^Ly6C^-^ cells) expressing CD206 (n=9) and **(B)** Ki67 (n=9). **(C)** Representative flow cytometry plots and summary graphs showing proportions and mean fluorescence intensity of CD45.1 host (resident) and CD45.2 donor (recruited) macrophages expressing Relmα following 4 days after i.p. injection of PBS or IL-4c at days 0 and 2 (n=4). For all experiments, data are shown as mean ± SEM and statistical comparisons were performed using paired *t*-tests. *p<0.05; **p<0.01; ***p<0.001; ****p<0.0001.

We investigated whether uterine macrophage subsets were functionally similar by systemic administration of IL-4c to shield-irradiated chimeric mice. Whilst both tissue resident and recruited macrophages responded readily, a greater proportion of recruited macrophages expressed the alternative activation signature molecule Relmα and recruited cells expressed higher levels of Relmα (Fig. 4C). Taken with evidence of a direct role for Relmα in tissue remodelling^40,41^, these data suggest that recruitment of monocytes and subsequent differentiation into macrophages may have important implications for function in the uterus.

### Properties of uterine monocytes and macrophages are conserved in humans and demonstrate differential responses to microbial stimulation

In healthy tissue, regulatory macrophages that maintain immune and tissue homeostasis are common, whereas pro-inflammatory monocytes and macrophages are predominantly associated with infection and inflammation^42^. During the reproductive (estrous) cycle in mice), we have demonstrated that dynamic infiltration of monocytes and subsequent differentiation into macrophages occurs in the absence of infection as part of normal tissue homeostasis (Figs. 2 and 3). To determine whether specific properties of uterine monocytes and macrophages are conserved in humans, we next characterised monocytes and macrophages from human endometrial biopsies. Similar to mice, we found that monocytes and macrophages in the uterus can be identified from a heterogenous pool of CD64^+^ cells. These cells did not express exclusion markers CD3 (T cells), CD19, CD20 (B cells) or CD66b (granulocytes), but did express HLA-DR. As uterine macrophage populations in mice were dependent on recruitment of CCR2^+^ monocytes (Fig. 3B), and CCR2^+^ cells in other mucosal tissues are recently infiltrating monocytes^36^, we differentiated monocytes from macrophages in human endometrial tissue as CCR2^+^ monocytes and CCR2^-^ macrophages (Fig. 5A). CCR2^-^ macrophages predominantly expressed CD163 which is associated with anti-inflammatory macrophages at other mucosal sites^17^. Akin to macrophages in the mouse uterus (Fig. 1), human endometrial macrophages nearly all expressed the alternative activation marker CD206 but very few expressed CD11c. Both monocytes and macrophages expressed CD14 but interestingly, although monocytes expressed CD11c as in mice, the majority of human monocytes in the uterus also expressed CD206 (Fig. 5A). This is important as studies characterising uterine macrophages in human tissue in terms of CD206 expression may also be including monocytes. These data indicate that monocyte and macrophage phenotypes in the uterus are conserved between mice and humans, including CCR2 being a feature of monocytes, high levels of expression of CD206 and limited expression of CD11c on macrophages.

**Figure 5:**
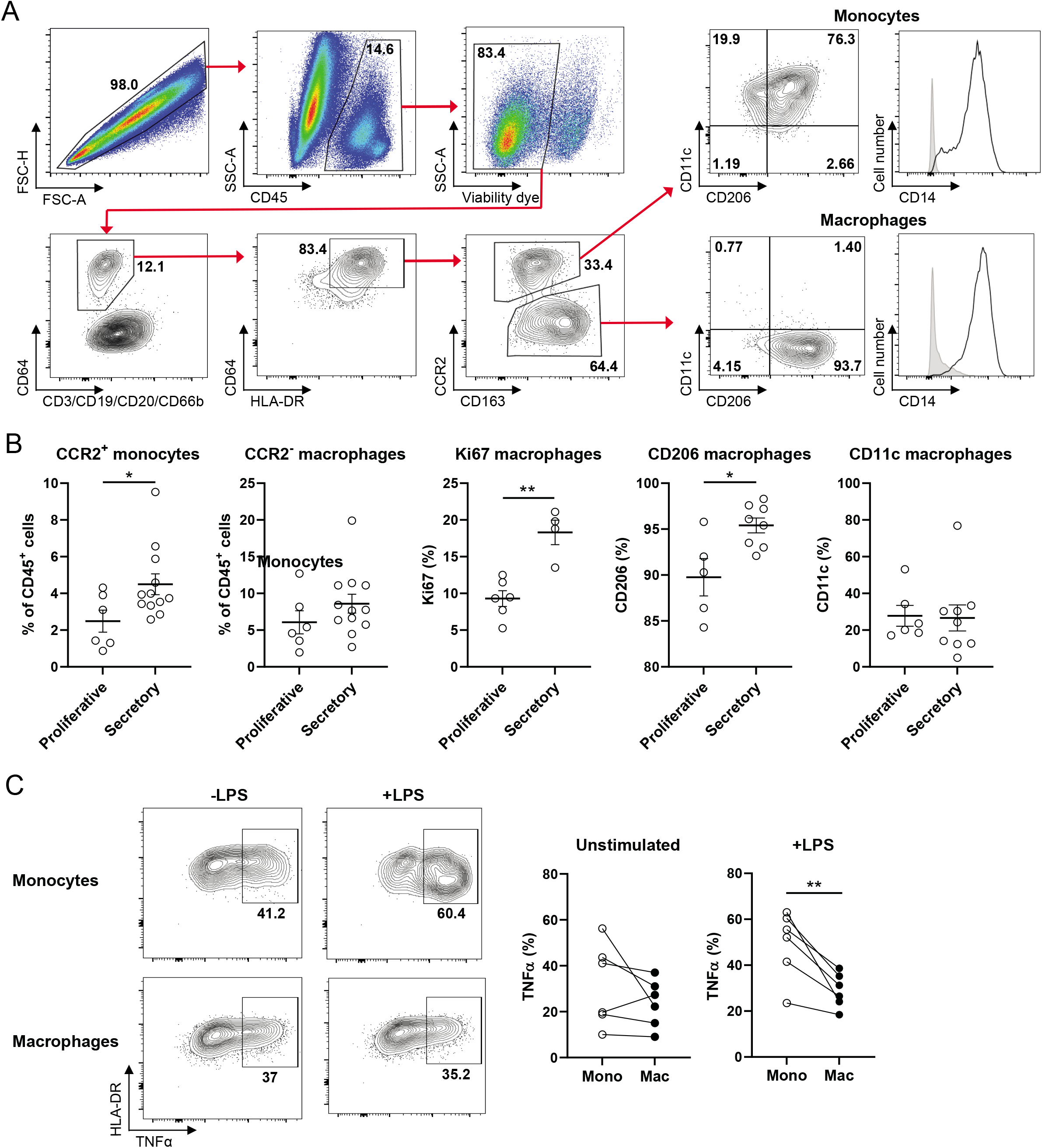
Characterisation of uterine macrophages in humans. **(A)** Representative flow cytometry plots depicting gating strategy for identification of monocytes and macrophages from human endometrial biopsies, and examples of expression of CD11c, CD206 and CD14 on monocytes and macrophages. Graphs are representative of >5 independent experiments (individual patients). **(B)** Summary graphs showing proportions of monocytes and macrophages of total CD45^+^ cells, Ki67 expression in macrophages, and CD11c and CD206 expression on macrophages from patients in proliferative (monocyte and macrophage proportions: n=6; CD11c: n=6; CD206: n=5; Ki67 n=6) and secretory (monocyte and macrophages proportions: n=12; CD11c: n=9; CD206: n=8; Ki67:n=4) phases of the menstrual cycle. Data are representative of >4 individual experiments, shown as mean ± SEM and statistical analyses were carried out using unpaired *t*-tests. **(C)** Representative flow cytometry plots and summary graphs showing proportions of monocytes and macrophages producing TNFα following 4 hour incubation +/- LPS (n=6). Data are representative of >4 individual experiments, shown as mean ± SEM and statistical analyses were carried out using paired *t*-tests. *p<0.05; **p<0.01.

We next assessed how monocyte and macrophage properties fluctuate throughout the human menstrual cycle. The proliferative phase corresponds to the proestrus phase of the estrous cycle in rodents, prior to ovulation where estrogen levels peak^43,44^, with lower levels of estrogen characteristic of the secretory phase in humans following ovulation when estrogen rapidly declines, akin to estrus in mice^44^. Studies have previously suggested a progressive increase in numbers of macrophages during the secretory phase of the cycle, with a peak at menstruation^45^, but these data were based on the marker CD68 which identified monocytes as well as macrophages^46^. We show here that similar to the increased numbers of monocytes in the uterus of mice during estrus (Fig. 2A), CCR2^+^ monocytes are increased in the secretory phase of the human menstrual cycle, following ovulation. As in mice, we observed no significant differences in proportions of macrophages among cycle stages (Fig. 5B). Nonetheless, macrophages from the proliferative and secretory phases of the menstrual cycle exhibited marked differences in Ki67 and CD206, potentially indicating increased proliferation of macrophages in the secretory phase of the cycle, and increased polarization towards alternative activation, respectively (Fig. 5B). In mice, CD206^+^ macrophages are essential for successful embryo implantantion^47^ supporting a role for these cells in regulation of tissue integrity in the uterus.

These data indicate the importance of characterizing monocytes and macrophages separately in the uterus to understand their separate dynamics and show that as in mice, monocytes are dynamic in their fluctuations in numbers whilst macrophages remain stable in number but exhibit cyclical differences in other properties including proliferation (Ki67). The fluctuations in monocyte and macrophage properties throughout reproductive cycles suggests that these events may be under hormonal control. Indeed, the functional remodelling that occurs in the endometrium throughout the menstrual cycle to enable support of a prospective pregnancy occurs in response to estrogen and progesterone. Estrogen in particular modulates innate and adaptive immune responses^48^, with monocytes^49–51^ and macrophages^52–54^ directly expressing estrogen receptors α and β (ERs).

Given evidence that estrogen can limit lipopolysaccharide (LPS)-induced inflammatory responses in macrophages in other tissues^52,55–57^, and that monocytes and macrophages in mucosal tissues respond differentially to LPS stimulation^17^, we next assessed the human endometrial monocyte and macrophage response to LPS *ex vivo*. For consistency, these experiments were carried out using samples collected during the secretory phase with comparison between monocytes and macrophages from the same patients (paired data). TNFα is typically produced by classically activated macrophages in response to LPS or other microbial stimulation. Surprisingly, despite displaying several characteristics of alternative activation, both monocytes and macrophages from human endometrial tissue constitutively expressed TNFα in the absence of LPS. There were no differences in TNFα production between unstimulated monocytes and macrophages, however monocytes were markedly more responsive to LPS stimulation than macrophages, with a greater proportion of monocytes producing TNFα after stimulation compared with macrophages (Fig. 5C).

To conclude, we have demonstrated previously unappreciated specialized features of macrophages in the uterus, including their potent responsiveness to type 2 stimuli and features of alternative activation. We have elucidated that several properties of uterine macrophages as conserved between mice and humans, and have highlighted the importance of distinguishing between monocyte and macrophages in the uterus. This is particularly important in light of the fluctuations in monocyte but not macrophage numbers during reproductive cycles and their differential responsiveness to microbial stimulation. These fluctuations in monocyte numbers taken with the potency of microbial stimulation for monocytes specifically, have critical implications for response to infection. We demonstrate that the uterus is one of the most replenished tissues of the body in terms of macrophage differentiation from recruited monocytes, with functional differences between recruited versus tissue-resident macrophages, indicating that recruited macrophages may be more polarized towards alternative activation. Overall, these data have important implications not only for the role of monocytes and macrophages in the uterus for regulating tissue repair, regeneration and remodelling that is essential for embryo implantation and pregnancy, but also for understanding immune responses to infections at different phases of reproductive cycles.

## METHODS

### Mice

Wild-type (WT) C57BL/6 (Envigo) were maintained under specific pathogen-free conditions at the University of Manchester, UK. *Ccr2^-/-^* and CD45.1 C57BL/6 mice were maintained under specific pathogen-free conditions at the University of Manchester, UK. Age-matched female mice (8-12 weeks) were used in all experiments, approved by the University of Manchester Animal Welfare Ethical Review Board and performed under licenses issued by the U.K. Home Office. All experiments were carried out according to U.K. Home Office regulations. For work performed at Institut Pasteur, animal experiments were conducted in accordance with approval of protocol number 2016-0010 by the *Comité d’éthique en expérimentation animale Paris Centre et Sud* and the *Comités d’Ethique pour l’Expérimentation Animale* Institut Pasteur (the ethics committee for animal experimentation), in application of the European Directive 2010/63 EU.

### Human samples and ethics

Human endometrial biopsies were obtained according to the principles expressed in the Declaration of Helsinki and under local ethical guidelines, and approved by the North of Scotland Research Ethics Service (reference number 19/NS/0163). Samples were obtained from patients undergoing hysteroscopic investigation at St. Mary’s Hospital, Manchester, UK, with findings of a normal endometrium or polyps during hysteroscopy, with no inflammatory disorders of the uterus (for example, endometriosis, Asherman’s syndrome). All patients provided written, informed consent for the collection of tissue samples and subsequent analysis. Patient diagnosis was confirmed by established clinical and histological criteria; patient demographic and clinical information is listed in Supplementary Figure 1.

### Reproductive cycle staging

In mice, stages of the estrous cycle were assessed at the time of tissue harvest, based on the proportions of cell types observed in vaginal secretions as previously described^28^. In humans, patients undergoing hysteroscopy were characterised as being in the secretory or proliferative phases of the menstrual cycle based on their usual cycle length, date of their last menstrual period and the date of the procedure. Ovulation was estimated at 14 days before the start of their next predicted menstrual period, with samples taken after this date classified as secretory phase, and samples taken before ovulation as proliferative phase.

### Murine uterus digest

Both uterine horns were removed from mice and collected in RPMI-1640 (Thermofisher Scientific), minced using fine scissors and incubated for 40 minutes in 1.25mg/ml collagenase D (Sigma), 0.85mg/ml collagenase V (Sigma) and 1mg/ml dispase (Gibco, Invitrogen) in a shaking incubator at 37°C. Tissue was then filtered through a 100μM filter and cells were analysed by flow cytometry.

### Human endometrial biopsy digest

Endometrial pipelle biopsies were collected in 5ml PBS and centrifuged to separate tissue from PBS supernatant which was then removed. Tissue pellets were then resuspended in 0.5mg/ml collagenase VIII (Sigma), 0.65mg/ml collagenase D (Sigma), 1mg/ml dispase (Gibco, Invitrogen), and 0.015mg/ml DNAse (Sigma) and incubated for 2x 20 minutes in a shaking incubator at 37°C. After each incubation, tissue was filtered through a 100μM filter. Remaining tissue was incubated again and tissue suspensions pooled following digests. Cells were washed and red blood cell lysis was carried out using ddH2O for 10 seconds following immediately by 1/10 volume 10x PBS. Cell were then analysed by flow cytometry or used for *in vitro* stimulation assays.

### Flow cytometry

0.2-2 x 10^6^ cells were stained at 4°C in the dark, in PBS with 2% FCS and 2% EDTA (FACS buffer) with Fc block (Biolegend) for 25 minutes. Antibodies used are listed in Supplementary Figure 2. For surface staining, cells were fixed in 2% paraformaldehyde (Sigma-Aldrich) for 30 minutes, washed and resuspended in FACS buffer before acquisition. For intracellular staining, following incubation with antibodies for surface staining, cells were washed in FACS buffer then fixed and permeabilised (eBioscience intracellular staining kit) before incubation for 60 minutes with antibodies for intracellular proteins (room temperature). Cells were washed and resuspended in perm buffer (eBioscience) before acquisition. Cells were analysed using an LSR Fortessa cytometer (BD Biosciences) and FlowJo software (Treestar).

### Immunofluorescent microscopy

Uteri were fixed with 4% paraformaldehyde (PFA) for 1 hour and washed with PBS. Samples were then dehydrated in 30% sucrose for 24 hours, then cut transversally and embedded in optimal cutting temperature compound (OCT), frozen, and sectioned at 30 μm. Sections were blocked for 1 hour 3% bovine serum albumin (BSA) + 0.1% Triton X-100 + Goat serum (1:20) in PBS, then washed three times. Immunostaining was performed using F4/80, and anti-GFP in staining buffer (0.5% BSA + 0.1% Triton X-100 in PBS) overnight. Sections were washed and stained with phalloidin (1:350) and secondary antibodies (1:2000) in staining buffer for 4 hours. The, sections were washed and stained with 4’,6-diamidino-2-phenylindole (DAPI). Confocal images were acquired on a Leica SP8 confocal microscope.

### Response to IL-4

IL-4 complex (IL-4c; IL-4/mAb to IL-4) was used as previously described^24,58^. Recombinant murine IL-4 (Biolegend) was combined with rat IgG1 mAb to IL-4, 11B11 (BioXCell), at a 1:5 molecular weight ratio^59^. Mice were injected i.p. with 5μg of IL-4c or PBS on day 0 and day 2. In some experiments, 50μl IL-4c or PBS was administered i.n. on day 0 and day 2. Tissues were collected on day 4 after initial injection.

### Uterus shielded bone marrow chimeras

For shielded irradiation, 8-10 week old wild type female CD45.1 C57BL/6 mice were anaesthetized by i.p. administration of ketamine (80 mg/kg; Vetoquinol) and xylazine (8 mg/kg; Bayer). Mice were irradiated with their abdomens positioned beneath lead shielding, and received 11 Gy irradiation as a split dose. Mice were reconstituted by i.v. injection with 5-10 x 10^6^ donor BM cells from CD45.2 (wild-type or *Ccr2^-/-^*) congenic mice. Mice were maintained on 0.03% enrofloxacin (Bayer) supplied through drinking water 1 week before and 4 weeks after irradiation. After irradiation mice were housed in autoclaved cages with sterile drinking water, food, bedding and enrichment. Reconstitution was allowed to occur for a minimum of 8 weeks before analysis. For experiments involving reconstitution of mice with WT BM only (University of Manchester), chimerism of uterine and colonic macrophages was normalised to that of circulating monocytes. For experiments involving comparison of reconstitution of mice with *Ccr2*^+/+^ and *Ccr2^-/-^* BM (Institut Pasteur), chimerism of uterine macrophages was normalised to that of circulating eosinophils, since this was the only circulating immune cell population that demonstrated comparable levels of chimerism in the blood of mice reconstituted with *Ccr2*^+/+^ or *Ccr2^-/-^* BM (Supplementary Figure 3).

### Response to LPS

Following tissue digest, human endometrial cells were incubated for 4 hours with monensin and Brefeldin A (Sigma) in the presence or absence of 100 ng/ml LPS (Sigma). Monocytes and macrophages were then identified by flow cytometry and assessed for intracellular cytokine production following cell permeabilization and fixation (eBioscience intracellular staining kit).

### Statistical analyses

Statistical analyses were performed using GraphPad Prism version 8.0/9.0 for Mac. Statistical differences were tested and described in the figure legends. Normality tests were performed on all datasets. Groups were compared using an unpaired *t*-test (normal distribution) or Mann-Whitney test (failing normality testing) for comparison of two groups. Paired *t*-test (normal distribution) or Wilcoxon matched-pairs signed rank test (failing normality testing) was used to compare results (e.g. from different cells or tissues) from the same individual. ANOVA with Tukey post-hoc testing (normal distribution) or Kruskal-Wallis test with Dunn’s post-hoc testing (failing normality testing) was used for multiple group comparisons. In all cases, a *p*-value of ≤ 0.05 was considered statistically significant. *p<0.05; **p<0.01; ***p<0.001; ****p<0.0001.

## ACKNOWLEDGMENTS

We gratefully acknowledge the National Institute for Health Research (NIHR) Clinical Research Network (CRN; Greater Manchester Core Team), Clinical Research Facility at Manchester University NHS Foundation Trust and the study participants for supporting this study. We gratefully thank R. Akhand and the CRN Research Nurse team in Reproductive Medicine (St. Mary’s Hospital, Manchester University NHS Foundation Trust) for their assistance with patient recruitment, consent, collection of biopsies and management of clinical data. We also thank G. Howell and M. Jackson in the FBMH Flow Cytometry Core Facility (University of Manchester), members of the BSF at the University of Manchester for help with animal work and J. Allen (University of Manchester), J. Brosens (University of Warwick) and A. Mowat (University of Glasgow) for critical evaluation of the manuscript. This research was funded by The Wellcome Trust and The Royal Society (206206/Z/17/Z to E.R.M.). L.L.M. was part of the Pasteur-Paris University (PPU) International PhD Program, which received funding from the European Union’s Horizon 2020 research and innovation program under the Marie Sklodowska-Curie grant agreement no. 665807 and from the Labex Milieu Intérieur (ANR-10-LABX-69-01). M.A.I. is supported by funding from the Agence Nationale de la Recherché (French National Research Agency) ANR-17-CE17-0014 and ANR-19-CE15-0015.

## AUTHOR CONTRBUTIONS

N.A.S. and L.L.M. provided critical material, expertise, carried out experiments and analysis and edited the manuscript.

L.M., P.R., O.M., J.D.A., D.R.B. and M.A.I. performed experimental and conceptual design, provided critical expertise and/or edited the manuscript.

E.R.M. performed the experimental and conceptual design, carried out experiments, data acquisition, analysis and wrote the manuscript.

## DATA AND MATERIALS AVAILABILITY

All data associated with this study are present in the paper or Supplementary Materials.

## Notes

### Competing Interest Statement

The authors have declared no competing interest.

