## Supplementary figures for "Macrophages in the uterus are functionally specialised and continually replenished from the circulation"

**Supplementary Figure 1: Patient clinical data**

| Diagnosis | Hysteroscopy findings | Age | Ethnicity | Cycle stage |
|---|---|---|---|---|
| Subfertility | Normal | 39 | Asian: Chinese | Secretory |
| Fibroids | Not submucosal | 39 | Unknown | Secretory |
| Subfertility | Polyp | 31 | Asian | Secretory |
| Recurrent miscarriage | Normal | 36 | Unknown | Secretory |
| Recurrent miscarriage | Normal | 28 | White | Secretory |
| Subfertility | Normal | 29 | Mixed | Secretory |
| Subfertility | Normal | 38 | White: Other | Secretory |
| Subfertility | Normal | 34 | White: British | Secretory |
| Recurrent miscarriage | Normal | 24 | White: British | Secretory |
| Subfertility | Normal | 36 | White: British | Proliferative |
| Subfertility | Normal | 32 | Black: African | Proliferative |
| Recurrent miscarriage | Normal | 36 | White: British | Proliferative |
| Subfertility | Uterine septum | 38 | White: British | Secretory |
| Subfertility | Normal | 38 | White: British | Proliferative |
| Recurrent miscarriage | Ovarian polyp | 28 | Asian: Pakistani | Proliferative |
| Recurrent miscarriage | Normal | 35 | White: British | Proliferative |

**Supplementary Figure 2: Antibodies**

**A**

| **Antibody** | **Reactive in** | **Conjugate** | **Clone** | **Source** |
|---|---|---|---|---|
| CD45 | Mouse | BV510 | 30-F11 | Biolegend |
| CD11b | Mouse | BV605 | M1/70 | Biolegend |
| CD206 | Mouse | BV421 | C068C2 | Biolegend |
| CD64 | Mouse | PeCy7 | X54-5/7.1 | Biolegend |
| CD11c | Mouse | BV395 | HL3 | BD Horizon |
| Ly6C | Mouse | PerCP-Cy5.5 | HK1.4 | Biolegend |
| Ly6G | Mouse | BV510, PerCP-CY5.5 | 1A8 | Biolegend |
| MHC class II (I-A/I-E) | Mouse | AF700 | M5/114.15.2 | Biolegend |
| Siglec F | Mouse | BV421 | E50-2440 | BD Biosciences |
| F4/80 | Mouse | FITC, APC | BM8 | Thermofisher |
| CX3CR1 | Mouse | BV605 | SA011F11 | Biolegend |
| CSF-1R | Mouse | APC | AFS98 | Biolegend |
| IL-4R$\alpha$ | Mouse | APC | I015F8 | Biolegend |
| Relm$\alpha$ | Mouse | PE (secondary with anti rabbit xenon labelling kit) | | Peprotech |
| Ki67 | Mouse | e450 | B56 | BD Biosciences |
| CD45.1 | Mouse | FITC | A20 | Biolegend |
| CD45.2 | Mouse | APC | 104 | Biolegend |

**B**

| **Antibody** | **Reactive in** | **Conjugate** | **Clone** | **Source** |
|---|---|---|---|---|
| CD45 | Human | BV510 | 2D1 | Biolegend |
| CD3 | Human | e450 | UCHT1 | Thermofisher |
| CD19 | Human | e450 | SJ25C1 | Thermofisher |
| CD20 | Human | e450 | 2H7 | Thermofisher |
| CD66b | Human | Pacific Blue | G10F5 | Biolegend |
| CD11c | Human | AF700 | Bu15 | Biolegend |
| CD206 | Human | PerCPCy5.5 | 15-2 | Biolegend |
| CD14 | Human | PerCPCy5.5 | 63D3 | Biolegend |
| CD64 | Human | APC | 10.1 | Biolegend |
| HLA-DR | Human | BV605, BV785 | L234 | Biolegend |
| CCR2 | Human | FITC, PE | K036C2 | Biolegend |
| CD163 | Human | BUV395, PeCy7 | GHI/61 | Biolegend, BD Biosciences |
| TNF$\alpha$ | Human | BUV395 | Mab11 | BD Biosciences |
| Ki67 | Human | BV605 | Ki-67 | Biolegend |

### Supplementary Figure 3

**A**

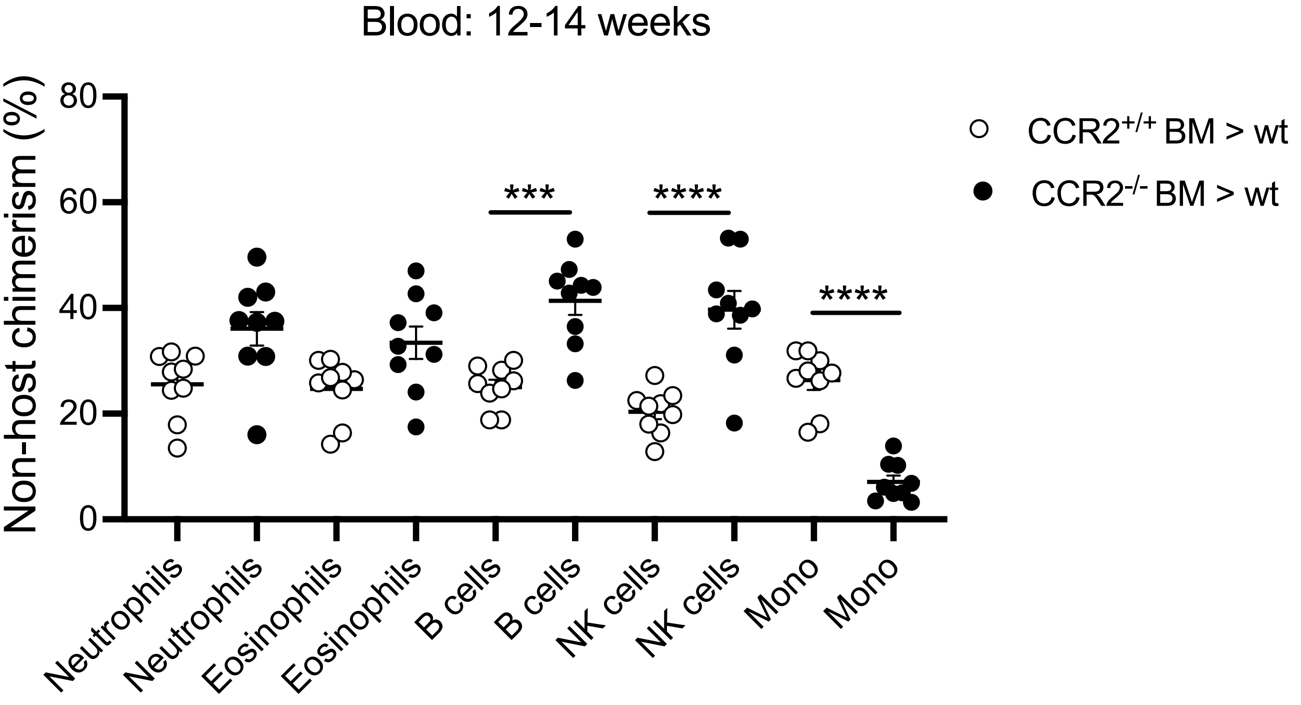


**B**

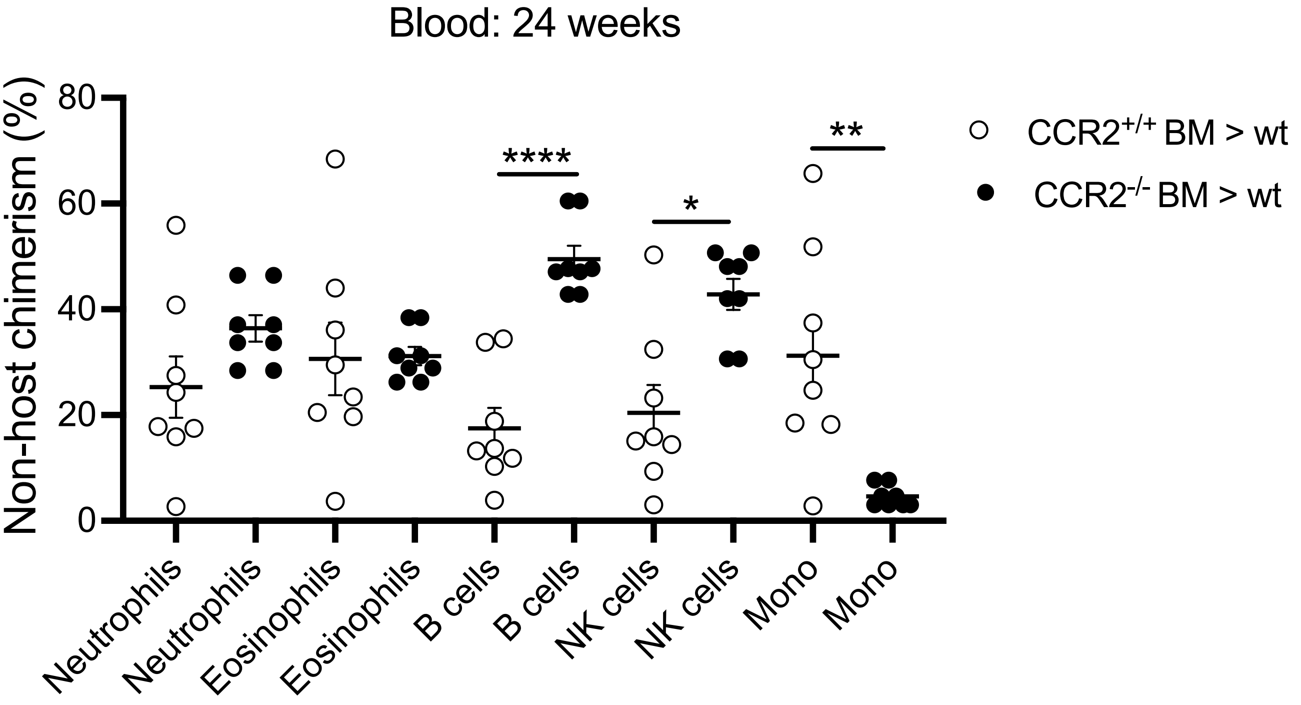

**Supplementary Figure 3: Chimerism in circulating immune cells.**

Shield-irradiated CD45.2 C57BL/6 mice were reconstituted with *Ccr2*$^{+/+}$ CD45.1 bone marrow (BM) and shield-irradiated CD45.1 C57BL/6 mice were reconstituted with CD45.2 *Ccr2*$^{-/-}$ BM. Data show the proportions of circulating immune cell populations reconstituted from *Ccr2*$^{+/+}$ and *Ccr2*$^{-/-}$ donor BM at **(A)** 12-14 weeks (n=9) and **(B)** 24 weeks (n=8). Data are representative of 2 independent experiments, shown as mean ± SEM and statistical comparisons were performed using 1-way ANOVA with Tukey post hoc corrections.
